# *Drosophila mef2* is essential for normal mushroom body and wing development

**DOI:** 10.1101/311845

**Authors:** Jill R. Crittenden, Efthimios M. C. Skoulakis, Elliott. S. Goldstein, Ronald L. Davis

## Abstract

MEF2 (myocyte enhancer factor 2) transcription factors are found in the brain and muscle of insects and vertebrates and are essential for the differentiation of multiple cell types. We show that in the fruitfly *Drosophila*, MEF2 is essential for normal development of wing veins, and for mushroom body formation in the brain. In embryos mutant for *D-mef2*, there was a striking reduction in the number of mushroom body neurons and their axon bundles were not detectable. D-MEF2 expression coincided with the formation of embryonic mushroom bodies and, in larvae, expression onset was confirmed to be in post-mitotic neurons. With a *D-mef2* point mutation that disrupts nuclear localization, we find that D-MEF2 is restricted to a subset of Kenyon cells that project to the α/β, and γ axonal lobes of the mushroom bodies, but not to those forming the α’/β’ lobes. Our findings that ancestral *mef2* is specifically important in dopamine-receptive neurons has broad implications for its function in mammalian neurocircuits.

## INTRODUCTION

Cellular identity is established during development by mechanisms that can be maintained for an animal’s lifetime and are conserved across evolution (Arendt, 2008; Arlotta & Hobert, 2015). Gene duplications can lead to functional variations among family members, thereby driving increased cell-type diversity (Arendt, 2008) and evolutionary pressure to maintain replicates (Assis & Bachtrog, 2013). To understand the most basic functions of a gene family it is expedient to evaluate the founding member. The MEF2 family of transcription factors has been assigned a myriad functions ranging from the differentiation of multiple cell lineages during development, to cellular stress response regulation and neuronal plasticity in adulthood. *D-mef2* is the single *Drosophila* homolog of the vertebrate *mef2a, b, c and d* family and as such can provide insight to conserved functions of this family.

*Drosophila mef2* and vertebrate *mef2* members exhibit considerable diversity in their transcriptional activation domains, but over 80% identity in the N-terminal sequences that encode the dimerization and DNA binding MEF and MADS domains (named for the evolutionarily conserved founding members MCM1, AGAMOUS, DEFICIENS, SRF) (Molkentin *et al.*, 1996; Potthoff & Olson, 2007). Correspondingly, the DNA sequences bound by MEF2 are evolutionarily conserved and MEF2 has been shown to activate transcription of orthologous gene sets in muscle, brain and immune systems of both flies and mice (Bour *et al.*, 1995; Lilly *et al.*, 1995; Ranganayakulu *et al.*, 1995; Lin *et al.*, 1997; Potthoff & Olson, 2007). The tissue specificity of MEF2’s actions is strongly influenced by the expression pattern of cofactors. Depending on which transcription factors MEF2 interacts with, immortalized cells in culture can be induced to display variable cell phenotypes: MEF2 and myogenin activate each other’s expression to initiate differentiation into skeletal muscle, MEF2 and Nkx2-5 activate each other’s expression to induce cardiac muscle formation, and MEF2 and MASH-1 activate each other’s expression to yield a neuronal phenotype (Skerjanc *et al.*, 1998; Ridgeway *et al.*, 2000; Skerjanc & Wilton, 2000). The specification and maintenance of cellular identity comprise major functions for MEF2. In both flies and mice, MEF2 is critical for the differentiation of multiple muscle cell lineages (Lilly *et al.*, 1995; Lin *et al.*, 1997; Potthoff & Olson, 2007). However, in mammalian neurons, a complex array of functions have been found for *mef2* family members in both development and neuroplasticity (Mao *et al.*, 1999; Okamoto *et al.*, 2000; Okamoto *et al.*, 2002; Flavell *et al.*, 2006; Shalizi *et al.*, 2006; Li *et al.*, 2008; Ryan *et al.*, 2013; Okamoto *et al.*, 2014; Chen *et al.*, 2016). The existence of different MEF2 family members within individual cells must be coordinated, and can even be antagonistic (Desjardins & Naya, 2017). Studies of a single ortholog, like the one in *Drosophila*, might serve to simplify this complexity and enhance our understanding of MEF2 by elucidating its most conserved functions.

*D-mef2* is expressed in *Drosophila* Kenyon neurons that make up the mushroom bodies (Schulz *et al.*, 1996), brain structures known for their functions in learning and memory (for review see Busto *et al.*, 2010 and Cognigni *et al.*, 2017). Mushroom bodies (MBs) are comprised of bilateral clusters of cell bodies that extend single neurites anteriorly to form a dendritic calyx and a fasciculated peduncle that branches to form axonal lobes. In adult *Drosophila*, each of the three major axon projection patterns, the branched α/β and α’/β’ lobes, or the γ axonal lobe (Crittenden *et al.*, 1998; Tanaka *et al.*, 2008) are formed from four neuroblasts that divide throughout embryonic, larval and pupal development (Lee *et al.*, 1999). The axonal lobes are segregated into domains according to their interconnections with distinct types of dopaminergic neurons and cholinergic mushroom body output neurons (Aso *et al.*, 2014). Numerous genes required for normal olfactory learning are preferentially expressed in the MBs, often in subsets of axonal lobes that likely reflects their distinct functions (McGuire *et al.*, 2001; Yu *et al.*, 2006; Krashes *et al.*, 2007; Akalal *et al.*, 2010; DasGupta *et al.*, 2014; Lim *et al.*, 2017). The role of *D-mef2* in MB development and function remains untested. Here, we examine the expression of D-MEF2 in developing mushroom bodies and among subsets of Kenyon cells in the adult fly, and evaluate mushroom body formation and overt phenotypes in *D-mef2* mutant alleles.

## MATERIALS AND METHODS

### *Drosophila* genetics

Fly stocks were raised at room temperature on standard sucrose and cornmeal media. The nine enhancer detector lines described were generated in a screen for mushroom body expression (Han *et al.*, 1996). The EMS, DEB, and γ ray mutants shown in Table 1 were identified in a screen for lethal genes at the cytological location 46C-F (Goldstein *et al.*, 2001). The parental chromosome for these lines was *adh cn pr* and they were maintained balanced over *CyO*.

**Table 1.** Intragenic complementation for lethality in *D-mef2* alleles. Eleven balanced *D-mef2* alleles were crossed inter se and the progeny counted. The viability is expressed as the percentage of trans-heterozygous progeny observed out of the number expected. The total number of progeny recovered is indicated in parenthesis.

| ***D-mef2***<br>**allele** | **22-21** | **22-24** | **25-34** | **26-6** | **26-7** | **26-49** | **30-5** | **44-5** | **48-7** | **66-65** |
| --- | --- | --- | --- | --- | --- | --- | --- | --- | --- | --- |
| **78-11** | 0% (192) | 0% (232) | 0% (217) | 0% (206) | 0% (0) | 1% (203) | 13% (97) | 0% (193) | 0% (220) | 0% (186) |
| **66-65** | 0% (221) | 0% (272) | 0% (260) | 0% (224) | 0% (191) | 0% (210) | 35% (194) | 0% (240) | 0% (266) |  |
| **48-7** | 0% (315) | 0% (307) | 0% (282) | 0% (230) | 0% (210) | 0% (203) | 0% (208) | 0% (304) |  |  |
| **44-5** | 0% (198) | 1% (251) | 6% (188) | 0% (246) | 0% (203) | 28% (307) | 14% (212) |  |  |  |
| **30-5** | 0% (210) | 3% (358) | 28% (207) | 0% (121) | 0% (128) | 78% (131) |  |  |  |  |
| **26-49** | 0% (174) | 0% (227) | 45% (252) | 0% (235) | 0% (234) |  |  |  |  |  |
| **26-7** | 0% (178) | 0% (406) | 0% (195) | 0% (195) |  |  |  |  |  |  |
| **26-6** | 0% (193) | 0% (291) | 0% (272) |  |  |  |  |  |  |  |
| **25-34** | 0% (223) | 0% (598) |  |  |  |  |  |  |  |  |
| **22-24** | 0% (519) |  |  |  |  |  |  |  |  |  |

### Molecular biology

Bacteriophage clones surrounding the enhancer detector insertion site in line 2487 were isolated from a Canton-S genomic library. The map constructed of the 46C region was expanded by 12 kb from coordinate 20 kb to 32 kb (Fig. 1) relative to the previously published maps (Bour *et al.*, 1995; Lilly *et al.*, 1995). The expansion was due to a stretch of repetitive DNA suggesting the likely insertion of a transposable element. Genomic DNA fragments adjacent to the insertions in lines 429, 883, 919, 2487, 3046, and 3775 were obtained by *Hind III* or *XhoI* plasmid rescue, according to previously described methods (Pirrotta, 1986). The insertion sites in lines 1484, 1828, and 2109 were determined by Southern blotting experiments.

**Figure. 1.**
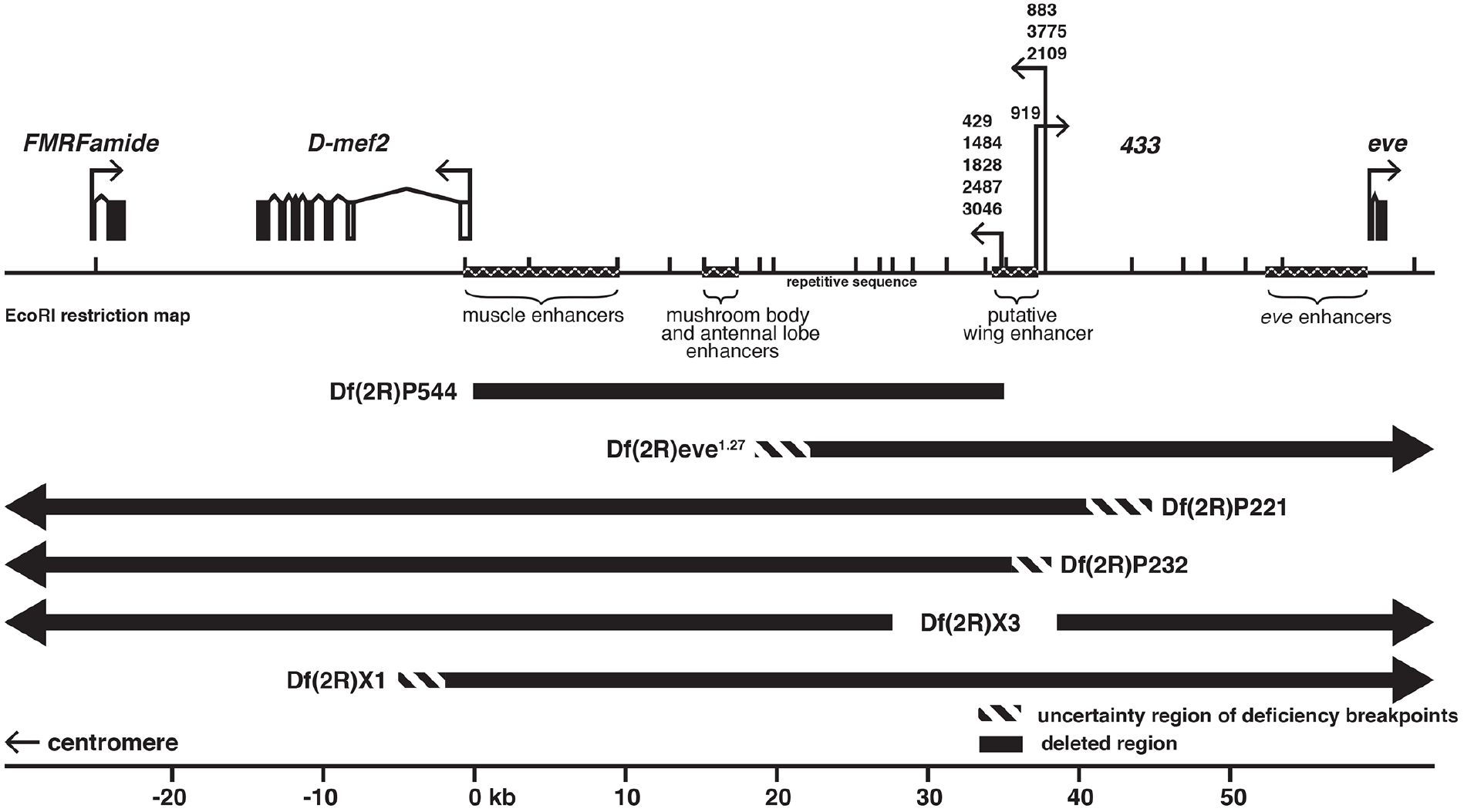
Enhancer detector lines reside distal to D-mef2. An EcoRI restriction map of the proximal region of 46C showing nine enhancer detector elements clustered upstream of the *D-mef2* gene. A single arrow represents insertions within 200 base pairs of each other; the direction of the arrow indicates the direction of *lacZ* transcription. The flanking gene *FMRFamide* is proximal to *D-mef2* and *even-skipped (eve)* is distal. The location of the mushroom body enhancer was derived from Schulz et al. 1996. Black bars below the map represent deficiencies in deletion lines. The open boxes of the *D-mef2* transcription unit represent untranslated exons and filled boxes represent exons in the open reading frame.

### Histology

β–galactosidase histochemistry and RNA *in situ* hybridization experiments were performed on frontal cryosections of the *Drosophila* head as previously described (Skoulakis & Davis, 1996). RNA probes were generated from the 5’ and the 3’ end of a *D-mef2* cDNA and used in separate experiments to validate results. Immunohistochemistry with chromogenic substrates was performed on paraffin and plastic sections of adult, larval, and embryonic *Drosophila* as previously described (Crittenden *et al.*, 1998). For immunofluorescence, FITC-or CY3-conjugated anti-rabbit and anti-mouse antibodies (Sigma) were used. Slides were mounted in Vectashield (Vector Laboratories). Antisera were raised against a fusion protein that contained both the MADS box and MEF domain of D-MEF2 and were provided by Drs. H. Nguyen and E. Olson.

### Cell counting experiments

We counted MB cells in embryos after immunofluorescent staining. Mushroom body cells were apparent in approximately 15 - 1 μm serial sections of each brain hemisphere. Each cell was visible in an average of 3.5 serial sections. Therefore, to estimate the number of MB cells per brain hemisphere, we divided the total number of cells counted by 3.5. Statistical comparisons between homozygous and heterozygous control embryos were made using unpaired, 2-tailed Student’s t-tests.

### Cell death

Acridine orange staining was performed as previously described (Abrams *et al.*, 1993). Homozygous *D-mef2* mutants were distinguished from sibling controls based on the absence of somatic muscles and the presence of bloated gut morphology.

### BrdU labeling

The treatment of larvae with BrdU to label dividing cells followed the protocol of Truman and Bate (Truman & Bate, 1988). For immunohistochemical detection of BrdU, paraffin sections of larvae were additionally treated with 2N HCl. For injections of embryos with BrdU, embryos were collected from timed egg lays on grape juice agar plates at 25 ^0^C. They were dechorionated according to standard procedures, aligned on double-sided tape, and covered with halocarbon oil. The posterior ends of the animals were then injected with 10 mg/ml of BrdU in insect Ringer’s solution (182 mM KCl, 46 mM NaCl, 3 mM CaCl_2_, 10 mM Tris-HCl, pH 7.2). The embryos were allowed to develop for three hours under halocarbon oil at 25 ^0^C and then fixed for 20 minutes in 50% Carnoy’s fixative in heptane with shaking. Finally, the embryos were sectioned in paraffin and prepared for immunohistochemistry following the protocol for larvae.

## RESULTS

### Enhancer detector lines identify distal *D-mef2* regulatory regions

From approximately 100 enhancer detector lines that were selected for mushroom body expression of the **β**–galactosidase reporter (Han *et al.*, 1996), we identified nine with insertions in cytological region 46C3 (Fig. 1). We mapped the transposon insertion sites by isolating plasmid rescue clones and using restriction mapping and DNA hybridization to compare to the 46C locus map (O’Brien *et al.*, 1994; Bour *et al.*, 1995; Lilly *et al.*, 1995). All nine lines harbored reporter gene insertions within a 3.5 kb region (Fig. 1) that is approximately 35 kb (including repetitive sequences) upstream of *D-mef2.*

The group of insertions more proximal to *D-mef2* (Fig. 1) presented preferential **β**–galactosidase activity in the MB and antennal lobes (Fig. 2). Additional expression throughout the cortex of the central brain and the optic lobes was exhibited by more distal insertions (Fig. 2). We compared the reporter **β**–galactosidase expression pattern to that of *D-mef2* mRNA and D-MEF2 protein in adult brain sections and found concordant enrichment in Kenyon cells and antennal lobe neurons (Fig. 3 A – F). These data suggest that reporter expression in the 46C enhancer detector lines is under the control of *D-mef2* MB and antennal lobe enhancers. A genomic fragment that is 17 kb proximal to the enhancer detector elements (Fig. 1) has been reported to possess MB enhancer activity (Schulz *et al.*, 1996). However, the deficiency *Df(2R)P544*, which was derived from enhancer detector line 2487 and lacks DNA sequence between *D-mef2* and the 2487 insertion site (Fig. 1), retained preferential β–galactosidase expression in the MB (not shown), suggesting that there are at least two mushroom body enhancer sequences at 46C.

**Figure. 2.**
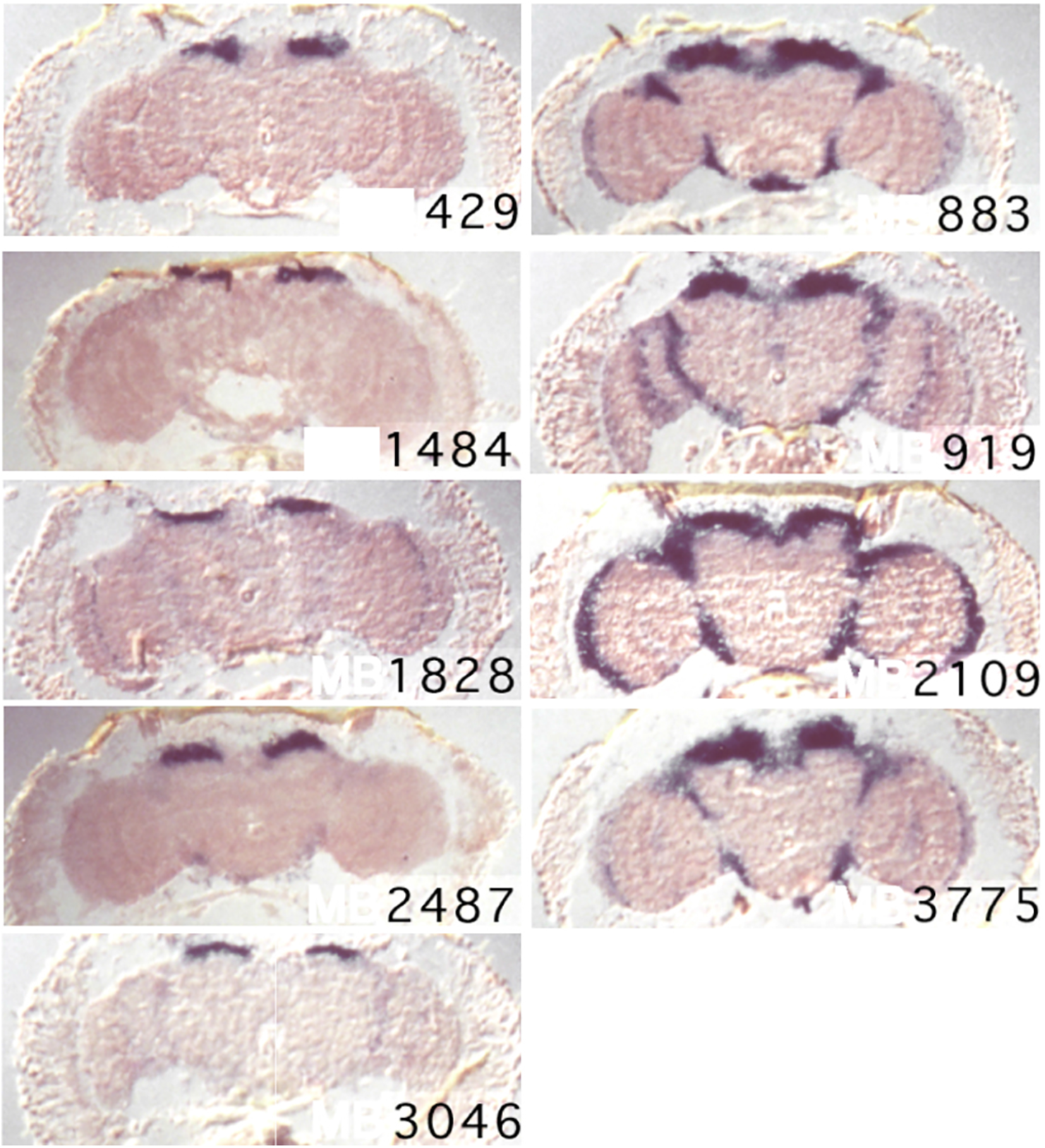
Enhancer detector lines reflect mushroom body and antennal lobe expression. Frontal sections of adult fly brains showing **β**–galactosidase activity in the mushroom bodies of nine enhancer detector lines with *lacZ* reporter gene insertions near *D-mef2.* Those lines with insertions proximal to the centromere (left column) exhibit **β**–galactosidase activity in the mushroom body nuclei whereas the lines with more distal insertions (right column) exhibit additional activity throughout the cortex (right column).

**Figure. 3.**
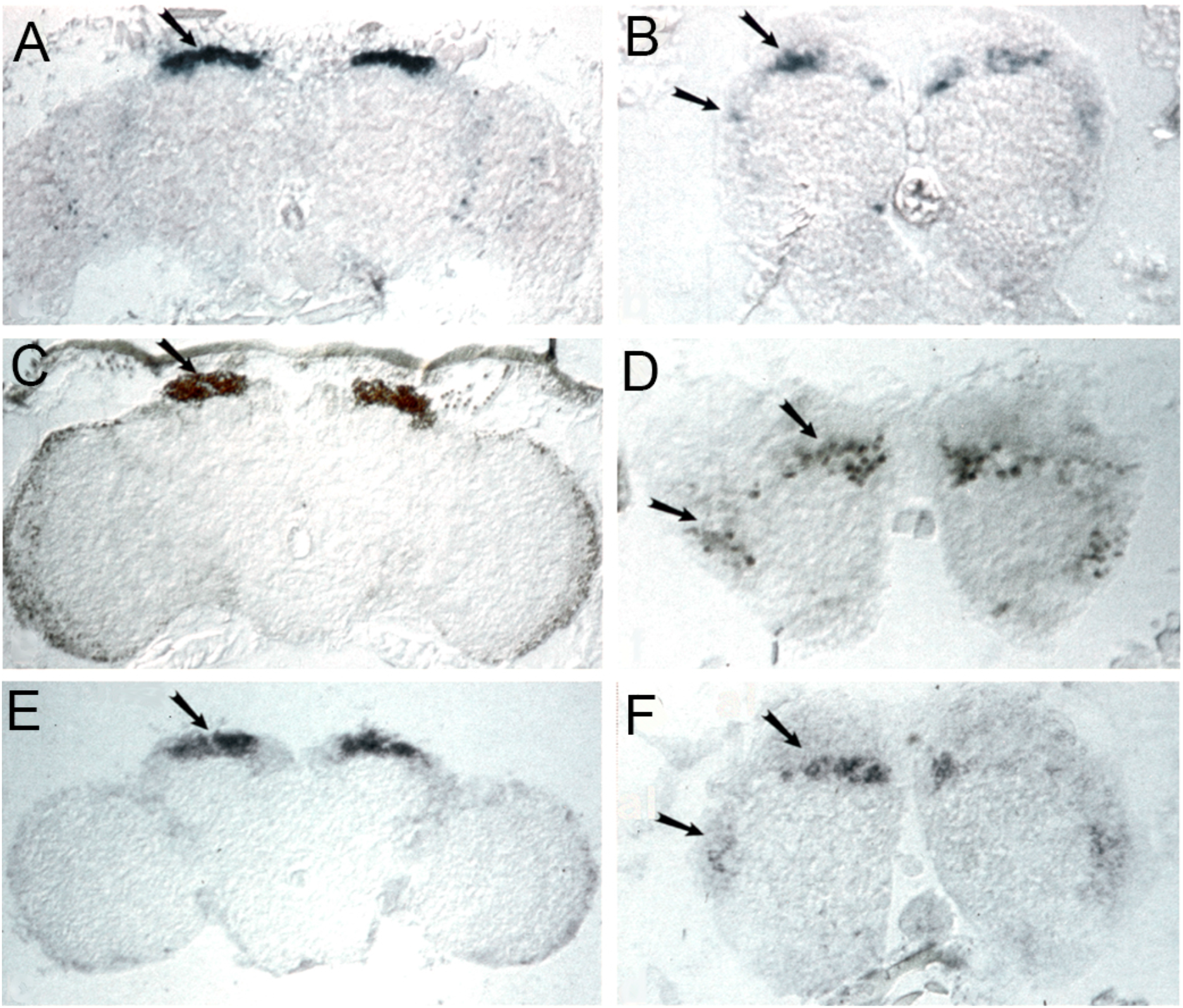
*D-mef2* mRNA and protein are enriched in adult mushroom body and antennal lobe neurons. Frontal sections through the adult brain showing the corresponding patterns of expression of the *lacZ* reporter gene, the *D-mef2* transcript, and D-MEF2 protein. Arrows in panels A, C, and E designate the mushroom body cells. Arrows in B, D, and F designate cells dorsal and lateral to the antennal lobe glomeruli. **A, B**) **β**–galactosidase activity in enhancer detector line 2487. **C, D**) Immunohistochemistry showing D-MEF2 protein distribution. **E, F**) RNA *in situ* hybridization showing *D-mef2* transcript distribution.

### *D-mef2* is required for normal wing development

Additional evidence that the enhancer detector lines are inserted within *D-mef2* regulatory sequences came from our discovery of a *D-mef2* wing phenotype. Enhancer detector line 919 showed strong expression and complete penetrance of disrupted wing morphology ranging from incomplete or ectopic cross-veins to bubbled wings (Fig. 4 A, B). A similar phenotype, at lower penetrance and expressivity, was observed in lines with insertions clustered more proximally to *D-mef2.* The wing phenotype was not observed in lines with P-element insertions more distal to *D-mef2.*

**Figure. 4.**
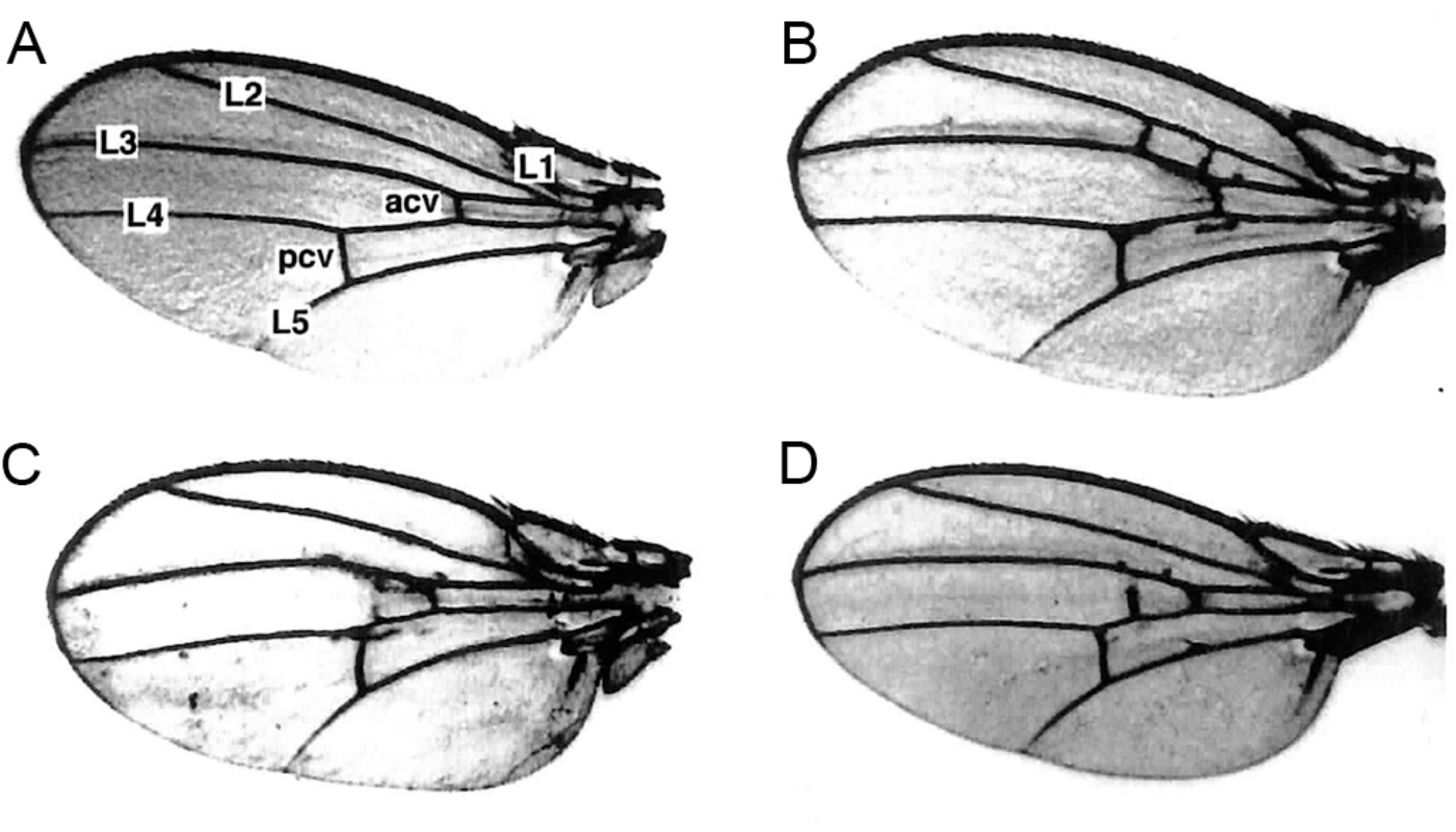
*D-mef2* is required for normal wing veination. **A**) A wildtype wing with veins labeled. acv: anterior cross-vein; pcv: posterior cross-vein; L1-L5: longitudinal veins. **B**) Ectopic veination in homozygous enhancer detector line 919. **C**) Ectopic veination in a transheterozygous *D-mef2^26-49/78-11^* fly. **D**) Non-complementation of the wing-phenotype in a transheterozygous*D-mef^22-21^*/Line 919 fly.

We observed a similar wing veination phenotype (Fig. 4 C) in flies with mutations that were determined to disrupt *D-mef2* function based on lack of complementation for viability with the deficiencies *Df(2R)X1* and *Df(2R)P520* (Bour *et al.*, 1995; Goldstein *et al.*, 2001). Six of these lines were generated by EMS (*D-mef2^22-21^, D-mef2^22-24^, D-mef2^25-34^, Dmef2^26-6^, D-mef2^26-7^*, and *D-mef2^26-49^*), three by DEB (*D-mef2^30-5^, D-mef2^44-5^*, and *D-mef2^48-7^*), and two by γ ray mutagenesis (*D-mef2^66-65^* and *D-mef2^78-11^).* By inter se complementation tests for viability (Table 1), we found that some alleles were strong (0% viability in combination), some medium (1 – 40% viability in any combination), and others weak (> 40% viability in any combination). In transheterozygous escapers, we frequently observed ectopic wing veination (e. g. Fig. 4 C).

To confirm that *D-mef2* dysfunction is responsible for the wing phenotype in the enhancer detector lines, we performed complementation tests with the null allele *D-mef2*^22-21^, which harbors a nonsense mutation in the 6th codon of *D-mef2* (Bour *et al.*, 1995). We observed wing blistering or abnormal veination in 74% of the transheterozygotes with line 919 (Fig. 4 D) and in 58% of transheterozygotes with line 429. Heterozygosity for P element insertion or *D-mef2^22-21^* alone did not cause a wing phenotype. Our results suggest that the *D-mef2*-proximal P-element insertions disrupt a wing enhancer, and establish a role for this protein in wing development.

### D-MEF2 is expressed in mushroom body neurons that send axonal projections into the α/β and γ lobes

We noticed that several clusters of mushroom body neurons lacked D-MEF2 immunoreactivity as determined by double-labeling (not shown) of wild-type animals with anti-LEONARDO (LEO), an immunomarker that exhibits global mushroom body expression (Skoulakis & Davis, 1996). D-MEF2-positive Kenyon cell subtypes were identified according to their axonal projection patterns, by evaluation of *D-mef2^26-49^* mutants that we discovered to harbor cytoplasmic D-MEF2. In *D-mef2^26-49^* mutants, anti-D-MEF2 decorated the axons of the α/β and γ lobe-projecting neurons but was absent from the α’/β’ lobes (Fig. 5 A – D). Although line *D-mef2^26-49^* was almost completely embryonic lethal, the few adult homozygous escapers (e. g. Fig. 5 A – D) showed grossly normal morphology and D-MEF2-immunoreactivity.

**Figure. 5.**
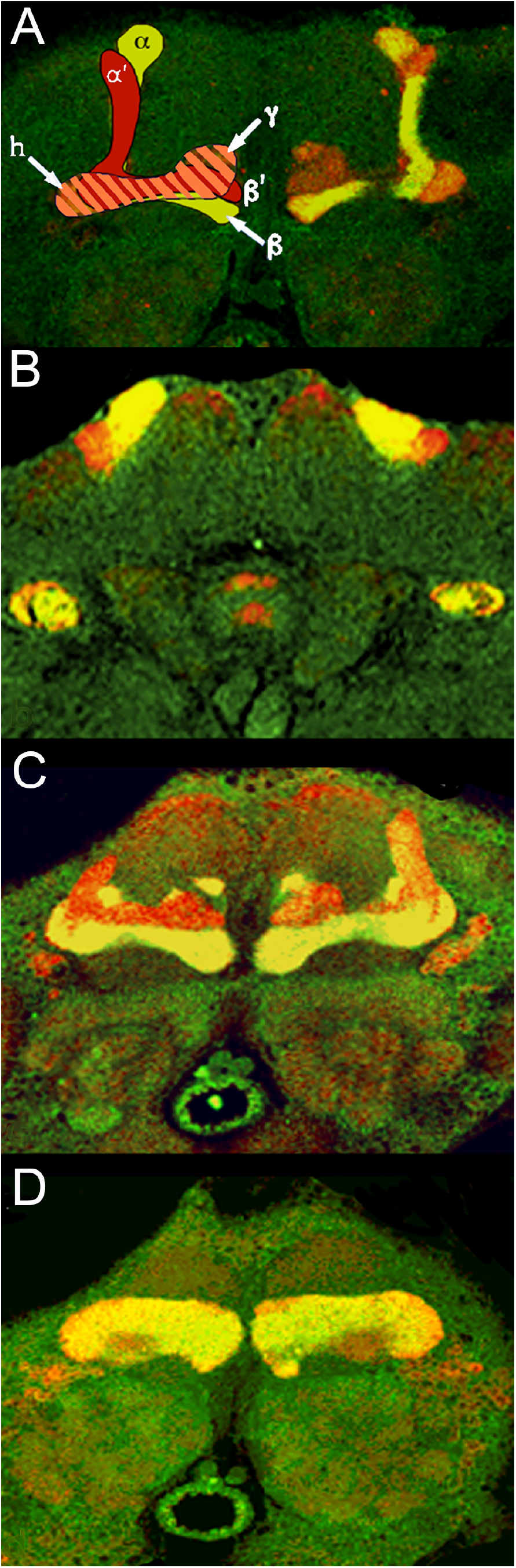
D-MEF2 is expressed in mushroom body neurons that project to the α, β and δ lobes, but not the α ‘ and β ‘ lobes. **A**) A frontal section through the central brain of a wild-type animal with an illustrative drawing over the lobes on one side. Mushroom body lobes were identified by gross anatomy and immunomarkers. Anti-LEO immunofluorescence (red) labels all five lobes whereas anti-FAS2 double-labeling (yellow) defines the core of the α/β lobe branches but not the α’/β’ lobes. The tips of the δ lobes, visible in this section, and the heel (h) also show light FAS2 immunoreactivity. **B – D**) In the *D-mef2^26-49^* line, frontal brain sections from posterior (B) to anterior (D) are co-immunolabeled for cytoplasmic D-MEF2 (green) and the pan-mushroom body marker LEO (red). Double-labeling is apparent in the α/β and δ lobes (yellow) whereas the α’/β’ lobes are not co-labeled for D-MEF2.

In horizontal brain sections from heterozygous *D-mef2^26-49^* mutants, D-MEF2 immunoreactivity was apparent in all four bundles of the posterior peduncle (Fig. 6 A, B), each of which is formed from the progeny of a single mushroom body neuroblast (Lee *et al.*, 1999). Thus, *D-mef2* is expressed in the descendants of all four mushroom body neuroblasts, but only those that project axons into the α/β branched lobes and into the γ lobes.

**Figure. 6.**
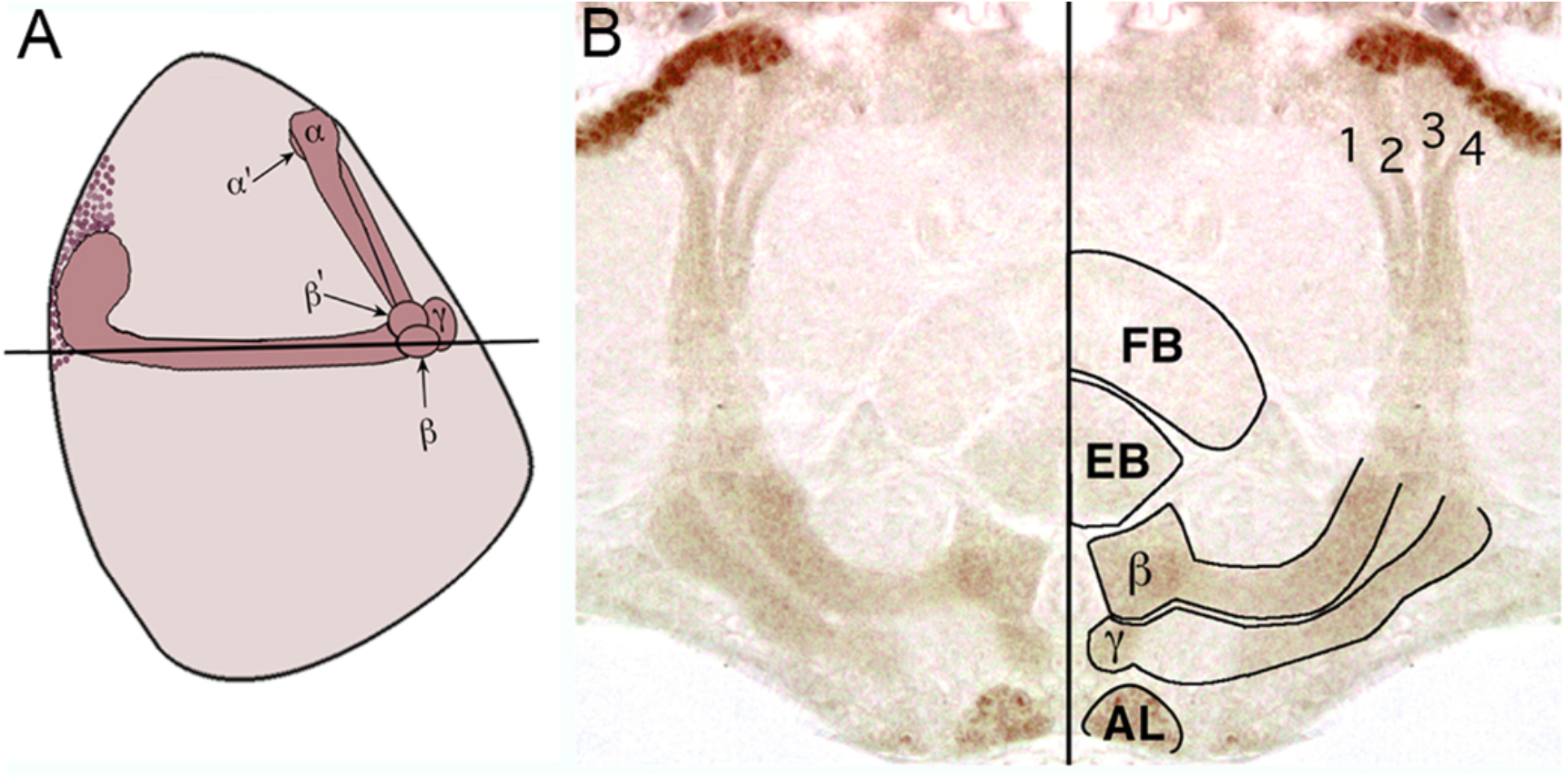
D-MEF2 is expressed in Kenyon cell descendants from all four mushroom body neuroblasts. **A**) A cartoon of the adult mushroom bodies in a sagittal plane, with anterior to the right. The black horizontal line represents the approximate plane of the horizontal section shown in (B). **B**) A horizontal section from a heterozygous *D-mef2^26-49^* adult shows D-MEF2 immunoreactivity (brown) in all four axon bundles of the posterior peduncle. In the mirrored image, the antennal lobe (AL), fan-shaped body (FB), and ellipsoid body (EB) are outlined and the four axon bundles arising from the Kenyon cells are numbered.

In the antennal lobe of *D-mef2^26-49^* flies, cytoplasmic D-MEF2 appeared restricted to the glomeruli and was not observed in projections of antennal lobe neurons or the antennal glomerular tract (Fig. 6 B), consistent with D-MEF2 expression in antennal interneurons. In the mutants, cytoplasmic D-MEF2 immunoreactivity was also detected in branches of the antennal nerve that extend into the antenno-mechanosensory center and into the antennal lobe (not shown), neurons that arise from the 2^nd^ and 3^rd^ antennal segments, respectively (Power, 1946). Correspondingly, nuclei within both antennal segments exhibited wildtype D-MEF2 immunoreactivity, a pattern also shown by the β-galactosidase expression in the enhancer detector lines (not shown). Other cells with D-MEF2 immunoreactivity in the head included muscles, photoreceptor cells, most cells of the lamina, and cells distributed throughout the medulla, lobula, and lobula plate.

### D-MEF2 is expressed in subsets of embryonic and larval mushroom body neurons

To explore the onset of *D-mef2* expression in the mushroom bodies, we surveyed expression from early stages of development. At embryonic stage 15, D-MEF2 was detectable in one or two cells in the dorso-posterior brain where mushroom body cells reside (Fig. 7 A). By stage 17, the number of brain cells expressing D-MEF2 in the mushroom body region had grown (Fig. 7 B), which is consistent with expression in a neuronal subtype that is dividing in late embryogenesis. Cell division appeared to be mostly restricted to the mushroom body region in late stage embryogenesis as evidenced by BrdU incorporation (Fig. 8). Moreover, in *D-mef2^26-49^* embryos, which display cytoplasmic D-MEF2 immunoreactivity, there was neuropil labeling in the central brain that resembled the mushroom body peduncle and dorsally oriented lobe (Fig. 7C). Double-labeling experiments with antibodies against D-MEF2 and against the Kenyon cell markers DACHSHUND (DAC) (Kurusu *et al.*, 2000; Martini & Davis, 2005) and against EYELESS (Kurusu *et al.*, 2000; Noveen *et al.*, 2000; Kunz *et al.*, 2012) showed only a partial overlap with D-MEF2 (not shown). We concluded that D-MEF2 is expressed in a subset of newly born Kenyon cells, from stage 15 to stage 17 of embryogenesis.

**Figure. 7.**
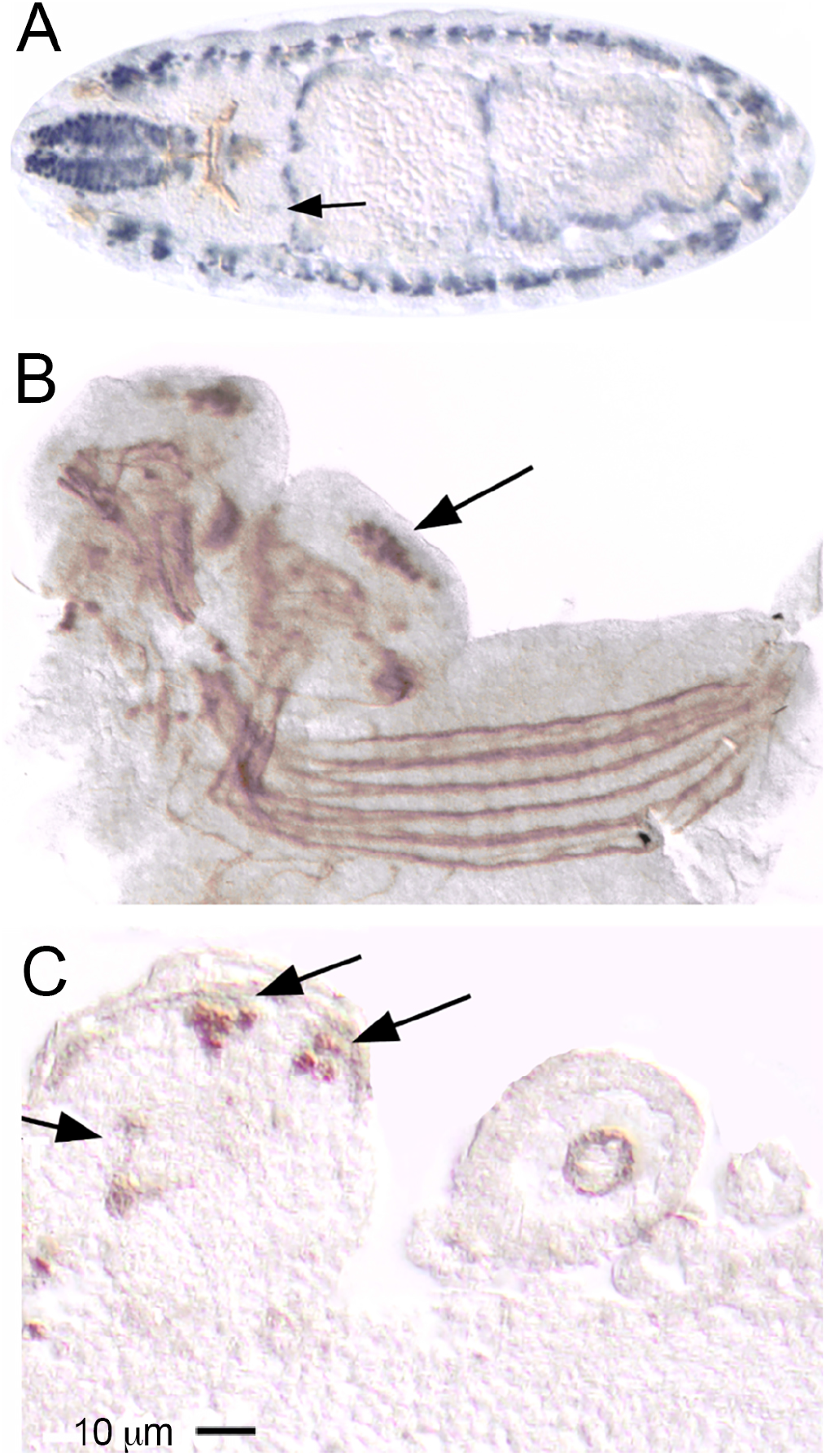
D-MEF2 is expressed in embryonic mushroom bodies. **A**) A horizontal section through a stage 15 wildtype embryo embedded in plastic and immunostained for D-MEF2 (alkaline phosphatase-coupled secondary antibody, blue) and FAS2 (horse radish peroxidase-coupled secondary antibody, brown). D-MEF2 expression is abundant in somatic and visceral muscle cell nuclei and is also visible in the dorso-posterior brain where mushroom body neurons are localized (arrow). FAS2 specifically labels axon tracts throughout the developing nervous system whereas D-MEF2 is localized to the nucleus. **B**) A whole-mount of the central nervous system dissected from a late stage 17 wildtype embryo and immunolabeled for D-MEF2 and FAS2 (both detected with horse radish peroxidase-coupled secondary antibody, brown). D-MEF2 expression is highly enriched in the mushroom body nuclei (arrow) and FAS2 is restricted to axons. **C**) A sagittal paraffin section through a late stage 17 embryo that is heterozygous for the *D-mef2^26-49^* mutation in which D-MEF2 is misexpressed in the cytoplasm. D-MEF2 immunoreactivity is apparent in the mushroom body nuclei (arrows in dorso-posterior brain), peduncle, and dorsally-projecting lobe (arrow). Anterior is to the left in A – C.

**Figure. 8.**
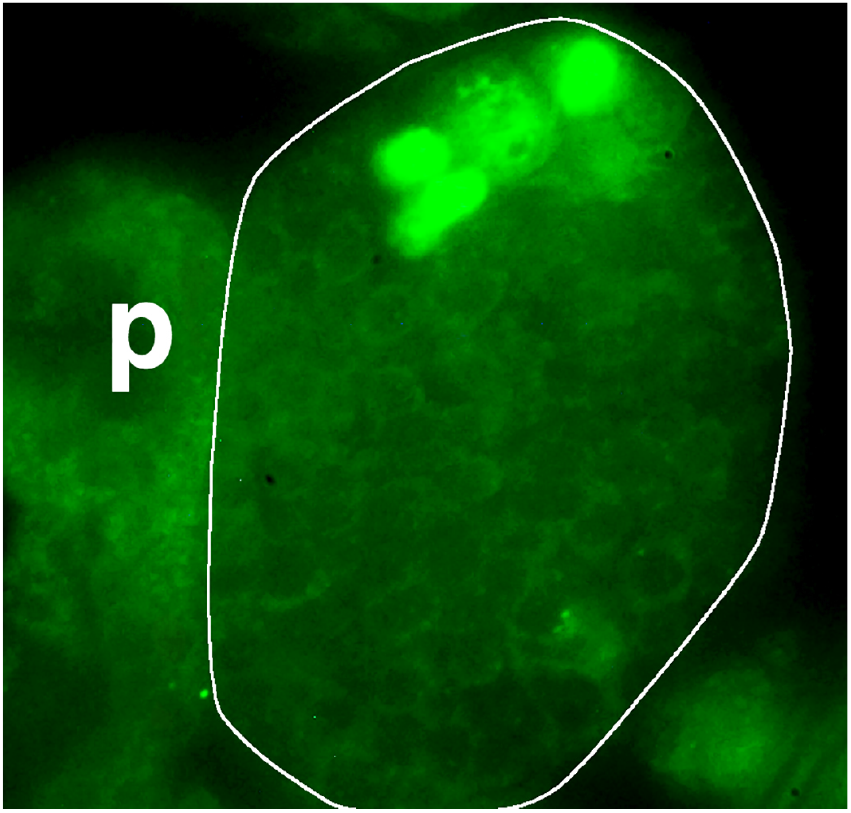
Cell proliferation is mainly restricted to mushroom body neurons in late stage embryogenesis. Anti-BrdU immunofluorescence (green) on a frontal hemi-section through the brain (outlined in white) of an embryo that was fixed 3 hours after BrdU was injected at 19 hours after egg laying. The pharynx (p) runs through the center of the brain. One mushroom body neuroblast, distinguished by its large size and diffuse labeling, is labeled in green along with three daughter cells.

At the first instar larval stage, D-MEF2 expression was confirmed to be in the post-mitotic Kenyon cells but not in the neuroblasts or ganglion mother precursor cells (Fig. 9 A, B). Weak D-MEF2 expression was also visible in cells surrounding, but not within, the single dividing neuroblast in the anterior brain (Fig. 9 A) that is known to give rise to a variety of antennal lobe cell types (Ito & Hotta, 1992; Stocker *et al.*, 1997; Lai *et al.*, 2008). In short, D-MEF2 was found in post-mitotic Kenyon cells and antennal lobe cells, but not in neuroblasts or ganglion mother cells of the developing brain.

**Figure. 9.**
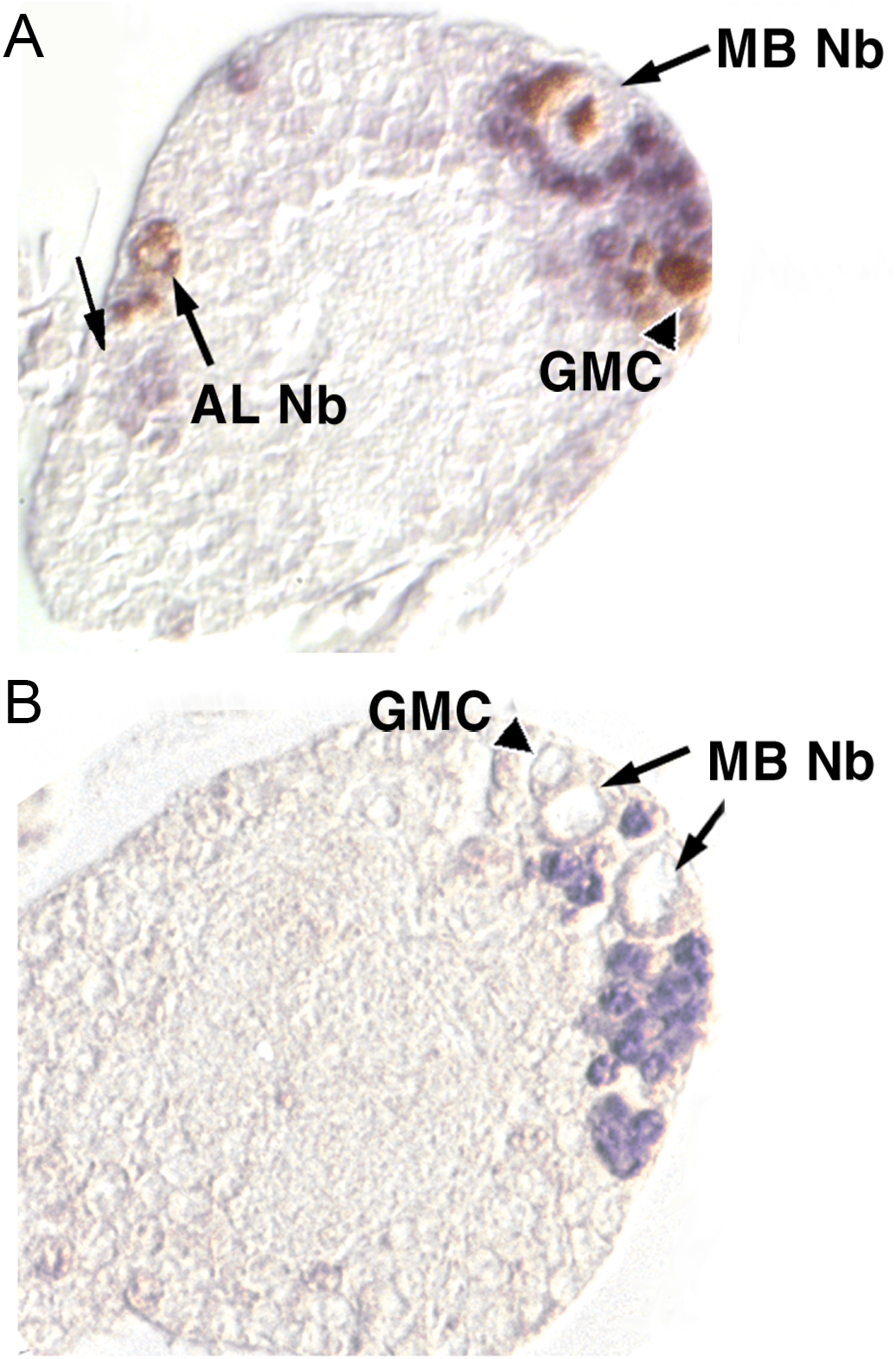
D-MEF2 is expressed in mushroom body neurons, but not their neuroblast or ganglion mother cell precursors. **A**) Sagittal sections through the brain of a first instar larva fed BrdU immediately after hatching and then immunolabeled for BrdU (brown) and D-MEF2 (blue); anterior is to the left. Anti-BrdU labels the mushroom body neuroblasts (MB Nb) and antennal lobe neuroblast (AL Nb) and their daughter cells (a putative ganglion mother cell, GMC arrowhead). Highly specific anti-D-MEF2 labeling is apparent in cells surrounding the mushroom body neuroblast, and more weakly staining cells are visible near the antennal lobe neuroblast (unlabeled arrow). **B**) A first instar larval brain immunolabeled only for D-MEF2 confirms the absence of D-MEF2 in neuroblasts and a putative ganglion mother cell.

### *D-mef2* is required for embryonic mushroom body formation

Considering that *D-mef2* was expressed in embryonic mushroom bodies, we tested for mushroom body malformation in severe *D-mef2* mutants that die as late stage embryos. We examined two different homozygous lethal lines as embryos, the null mutant *D-mef2*^22-21^, and *D-mef2^26-6^*, which carries a point mutation that disrupts the DNA binding domain of the protein, but retains its expression (Nguyen *et al.*, 2002). Although the heterozygous control and homozygous mutant embryos formed cuticle at the same time, gut distension was a prominent *D-mef2* mutant phenotype (Ranganayakulu *et al.*, 1995) in the homozygotes. Homozygotes were further distinguished by the absence of muscle immunolabeling for D-MEF2 in line *D-mef2^22-21^* and myosin heavy chain in line *D-mef2^26-6^* (Bour *et al.*, 1995; Lilly *et al.*, 1995).

We used the mushroom body immunomarker DAC to count mushroom body neurons in consecutive sagittal sections through stage 16 homozygous and heterozygous *D-mef2^22-21^* animals (Fig. 10 A). Anti-DAC immunoreactivity was observed in an estimated average of 63 cells per dorso-posterior brain hemisphere in the heterozygotes, compared to only 36 cells per hemisphere in the null homozygotes (Fig. 10 B), representing a 43% reduction in the number of DAC-positive mushroom body neurons. This loss was not consequent of failed neuroblast formation, as in the process of cell counting we observed four neuroblasts in each hemisphere of the *D-mef2^22-21^* homozygotes, and we also confirmed that these neuroblasts do incorporate BrdU upon injection 19 hours after egg laying (not shown).

**Figure. 10.**
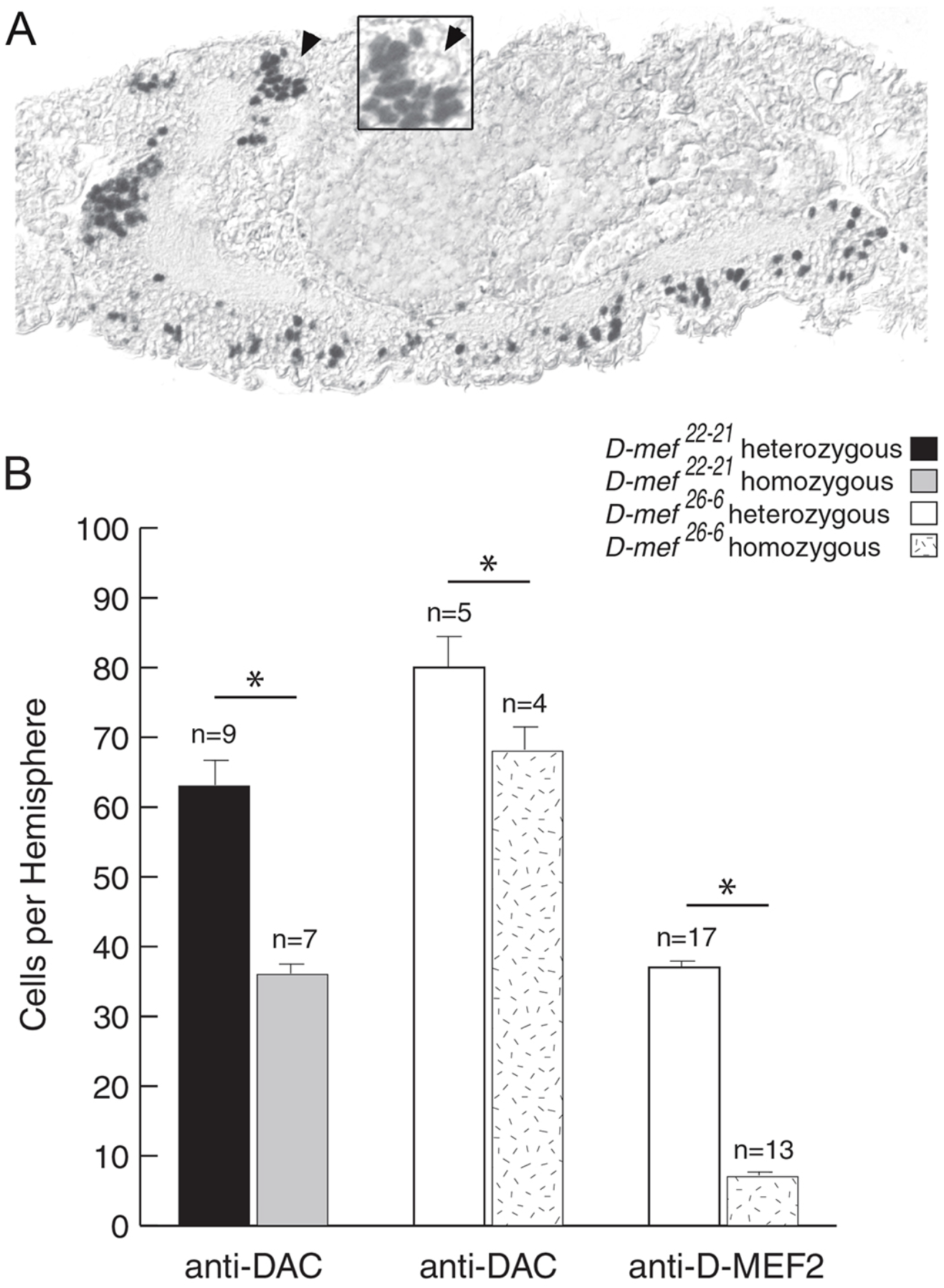
*D-mef2* mutant embryos have a paucity of mushroom body neurons. **A**) A sagittal section through a null mutant *D-mef2^22-21^* embryo immunostained for DAC (black) to label mushroom body neurons located in the dorso-posterior brain (magnified in inset). The mushroom body neuroblast is not labeled for DAC (arrowhead). **B**) Counts of mushroom body neurons that were immunolabeled for DAC or D-MEF2. *D-mef2^22-21^* and *D-mef2^26-6^* homozygous mutant embryos had significantly fewer mushroom body neurons than their age-matched heterozygous controls (P < 0.05 for a all three pair-wise comparisons by Student’s unpaired 2-tailed t-test). The number of brain hemispheres evaluated is indicated above the bars on the graph. Error bars show standard errors of the mean.

We made similar cell counts in stage 16 homozygous and heterozygous embryos from line *D-mef2^26-6^.* The control heterozygotes had an average of 80 DAC positive mushroom body neurons, whereas the homozygous mutants had an average of 68 (Fig. 10 B), representing a 15% reduction. We also counted the number of D-MEF2-positive neurons in this mutant. An average of 37 cells were counted per dorso-posterior hemisphere in the controls, whereas only 7 were found on average in the homozygous mutants (Fig. 10 B), an 81% reduction. Double-labeling experiments in controls showed that DAC was expressed in a greater proportion of mushroom body neurons than D-MEF2 (not shown). Therefore, a more substantial effect on D-MEF2-expressing Kenyon cells than on DAC-expressing neurons would be expected if the phenotype is cell autonomous as suggested by our observations. The difference in the number of DAC positive cells between the *D-mef2^22-21^* and *D-mef2^26-6^* heterozygous animals is likely to be due to a slight difference in the ages of the animals between experiments; however, since the heterozygous and homozygous animals within each genotype were aged and collected together, our primary findings were not compromised.

The inability of *D-mef2* mutants to undergo complete mushroom body formation was further indicated by their loss of mushroom body neuropil. We assessed neuropil immunolabeling with the embryonic mushroom body markers FAS2 and the protein kinase A subunit DCO (Skoulakis *et al.*, 1993; Crittenden *et al.*, 1998; Cheng *et al.*, 2001). In stage 17 heterozygous *D-mef2^22-21^* embryos, the immunostained peduncle and lobes (Fig. 11 A, C, E) appeared similar to what we had observed with these and other markers previously (Crittenden *et al.*, 1998). In contrast, neither anti-FAS2 nor anti-DCO labeled mushroom body structures in any sections from homozygous *D-mef2^22-21^* embryos processed on the same slides as controls (Fig. 11 B, D, F). It is unlikely that the failure to see immunostaining is due to a role for D-MEF2 in regulating the expression of both of these markers because ectopic D-MEF2 expression using the *GAL4/UAS* system with 5 different drivers failed to reveal ectopic expression of DCO or FAS2 (not shown). Furthermore, no reduction in DCO or FAS2 expression was evident in the brains of adult hypomorphic *D-mef2* mutants (not shown). In summary, we found that severe hypomorphic and null *D-mef2* mutants have reduced numbers of differentiated mushroom body neurons and a failure of mushroom body formation in embryogenesis.

**Figure. 11.**
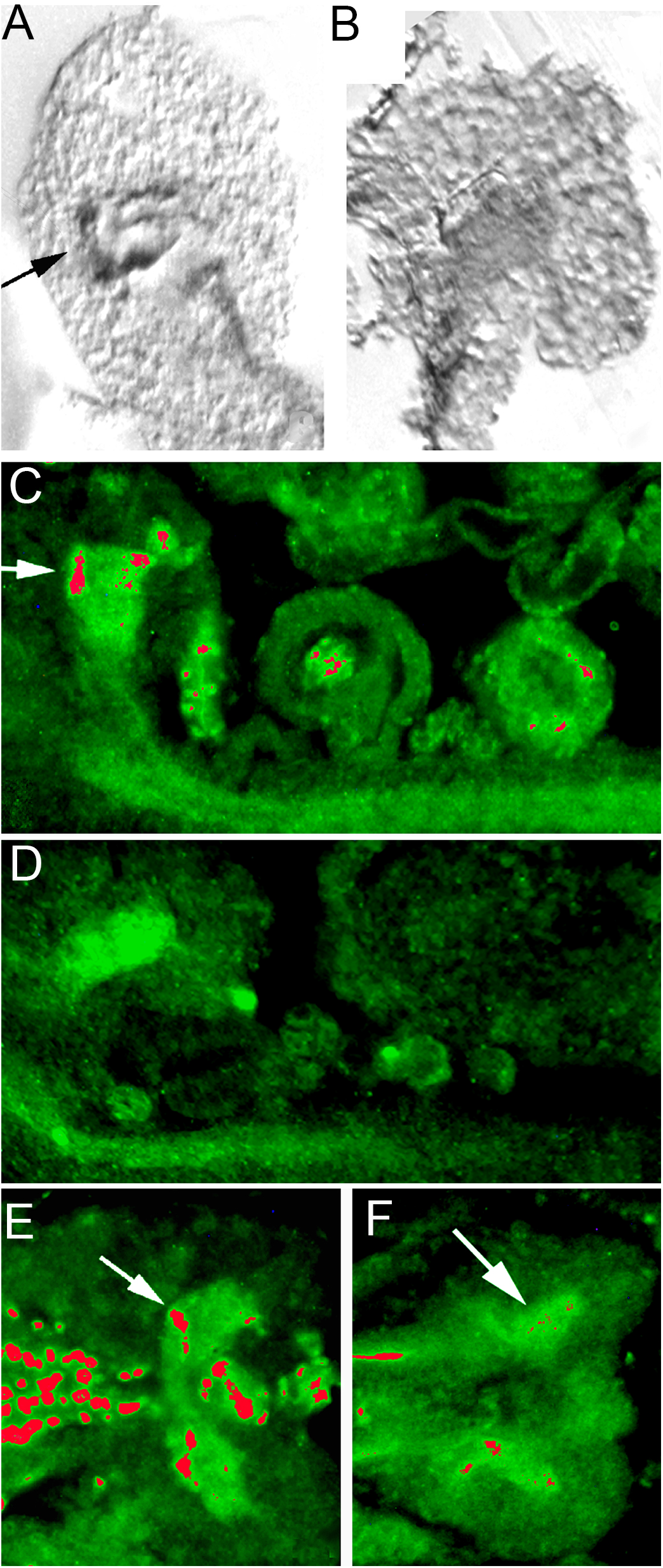
Mushroom bodies fail to form in *D-mef2* mutants. **A, B**) Anti-FAS2 immunohistochemistry on sagittal sections through the brains of stage 17 embryos; anterior is to the left. A) The mushroom body marker anti-FAS2 reveals the peduncle and the vertical mushroom body lobe (arrow) in heterozygous *D-mef2^22-21^* embryos; anti-FAS2 also labels the cervical connectives, but it does not label cell bodies. **B**) No mushroom body neuropil is detectable with anti-FAS2 in the brains of homozygous *D-mef2^22-21^* mutants. **C, D**) The mushroom body marker anti-DCO (green, with highest intensity false-colored in red) was exposed to sagittal sections through the central nervous system of stage 17 embryos that are heterozygous (A) or homozygous (B) for *D-mef2*^22-21^. Anterior is to the left. Anti-DCO decorates the central nervous system neuropil in both embryos but the mushroom body lobes (arrow in C) are only labeled in the control. Anti-D-MEF2 (green) labels the nuclei of muscles and mushroom body neurons in controls and was included for genotyping purposes. **E, F**) Symmetrical horizontal sections through the brain labeled with anti-DC0 and anti-D-MEF2 (green immunofluorescence). Anterior is to the left. E) In the center of the photograph, anti-DC0 reveals the central brain neuropil of the two brain hemispheres in this control embryo. The arrow points to the medially extending lobe. Pharyngeal muscle nuclei on the left are strongly labeled for anti-D-MEF2. F) Anti-DC0 exposed to *D-mef2* null embryo sections shows general neuropil labeling but does not reveal any mushroom bodies. An arrow points to where the lobes should be observable. D-MEF2 is not detectable in the muscles, confirming the genotype of these animals.

## DISCUSSION

### *D-mef2* functions in wing veination

P element insertions upstream of *D-mef2* led to our discovery of a wing veination function for *D-mef2.* The 46C enhancer detector lines did not show overt myogenesis or mushroom body development problems but did show ectopic wing veination or bubbling that is non-complementary with *D-mef2* mutations and that appears identical to what we found in viable transheterozygous *D-mef2* point mutants. Overexpression of *D-mef2* was found, in a large-scale screen of transcription factors, to induce wing blistering (Schertel *et al.*, 2015) but it was not investigated further. Screens for wing veination phenotypes have identified over 300 genes with enrichment for members of the Notch, EGFR and Dpp (TGF-β homolog) signaling pathways, considered essential for intercellular communication (Molnar *et al.*, 2006; Bilousov *et al.*, 2014). MEF2 can be linked to the regulation of these pathways – for example, *Tkv (thick veins)*, which encodes a Dpp receptor, is repressed by D-MEF2 during *Drosophila* egg formation (Mantrova *et al.*, 1999). Indeed, disruptions in *Dpp* and *tkv* expression can result in anterior cross-vein and blistering phenotypes that are similar to what we found in *D-mef2* hypomorphs (de Celis, 1997). Another member of the Dpp-Tkv pathway is p38 mitogen-activated protein kinase, which phosphorylates and activates mammalian MEF2 (Mao *et al.*, 1999) and in its dominant negative form causes ectopic wing veination in flies (Adachi-Yamada *et al.*, 1999). Collectively with our results, these data suggest that the abnormal vein formation in hypomorphic *D-mef2* mutants is caused by a failure in the Dpp-Tkv pathway.

### Nuclear retention signal for D-MEF2

Mammalian MEF2 contains several sequences near the C-terminus that are required for its nuclear localization, but these sequences are not conserved in *Drosophila* and the D-MEF2 nuclear localization sequence has not been identified (Yu, 1996; Borghi *et al.*, 2001). We identified a mutant, *D-mef2^26-49^*, in which D-MEF2 fails to be retained in the nucleus. The mutation in line *D-mef2^26-49^* was previously described as a missense point mutation that converts amino acid 148 from Thr to Ala (Lovato *et al.*, 2009). From a BLAST^®^ comparison to mouse MEF2 it appeared that this Thr is conserved in MEF2A but not in other MEF2 family members. This region of the protein is evolutionarily conserved and is termed the HJURP-C domain (Holliday junction regulator protein family C-terminal repeat). The HJURP-C domain is present in MEF2A, MEF2C and MEF2D but is lacking in MEF2B. The function of the HJURP-C domain is poorly understood but our results suggest that it is required for nuclear localization of D-MEF2.

### Mushroom body expression pattern of *D-mef2*

Previous reports have shown that mushroom body neurons begin to differentiate at stage 14 and continue to be born until shortly before pupal eclosion (Ito & Hotta, 1992). Our embryonic expression studies indicated that D-MEF2 becomes detectable in the mushroom body neurons as early as stage 15. In the embryo and larva, D-MEF2 immunoreactivity was in post-mitotic Kenyon cells and antennal lobe neurons, but not in neuroblasts or ganglion mother cells, consistent with the developmental expression profile in the honeybee *Apis mellifera* (Farris *et al.*, 1999). Likewise in mammals the initiation of *mef2* expression in cortical neurons coincides with their exit from the cell cycle (Lyons *et al.*, 1995; Mao *et al.*, 1999). Thus, the expression profile of *mef2* is consistent with a role in specification of neuronal cell identity in both insects and mammals during development.

Mushroom body neurons that give rise to the different lobes are generated sequentially from the four dorsal posterior neuroblasts and are interdependent for pathfinding and survival (Kurusu *et al.*, 2002; Martini & Davis, 2005). In adults, we found D-MEF2 expression in all four tracts of the posterior peduncle, indicating D-MEF2 expression in descendants of all four mushroom body neuroblasts. However, based on double-labeling experiments with other Kenyon cell markers, D-MEF2 is expressed in only a subset of mushroom body neurons in the embryonic and adult stages. The cytoplasmic mislocalization of D-MEF2 in line *D-mef2^26-49^* served to show that D-MEF2 is expressed at high levels in Kenyon cells that form medially and vertically-extending lobes in the embryo and in adult, α/β- and γ-lobe forming neurons, but not in the α’/β’ neurons. Mutant cytoplasmic D-MEF2 showed that the antennal lobe expression appeared to be confined to interneurons whereas projection neurons were found in the antennal segments that house olfactory receptors, hygroreceptors, thermoreceptors and the sound-sensing Johnston’s organ (Stocker, 1994). These structures are serially linked in the pathway for odor perception (Power, 1946): odor detection occurs in olfactory neurons of the third antennal segment, which synapse onto projection neurons in the antennal lobe glomeruli that in turn send sensory information to the mushroom body calyces. Mammalian MEF2 expression is maintained into adulthood and is important for neuronal plasticity (Flavell *et al.*, 2006; Sivachenko *et al.*, 2013; Chen *et al.*, 2016; Chen *et al.*, 2017). Together with the D-MEF2 expression pattern that we found, these data suggest that D-MEF2 is important for plasticity differences between the mushroom body lobes (Yu *et al.*, 2006) in olfactory learning.

MEF2 interacts physically with myogenic and neurogenic factors to potentiate cell-type specific gene transcription (Molkentin *et al.*, 1995; Black *et al.*, 1996; Mao & Nadal-Ginard, 1996). The D-MEF2 mushroom body lobe expression pattern expression gives clues to possible transcriptional interactors for D-MEF2. Examples of mushroom body markers with similar Kenyon cell subtype distribution to D-MEF2 include FOXP (DasGupta *et al.*, 2014), HDAC4 (Fitzsimons *et al.*, 2013), DRK (Crittenden *et al.*, 1998; Kotoula *et al.*, 2017), and FAS2 (Crittenden *et al.*, 1998; Cheng *et al.*, 2001). MEF2 interactions with several of these molecules have already been established. In mammals, HDAC4 (histone deacetylase 4) is known to bind to MEF2 to repress transcription and *Drosophila* HDAC4 is important for muscle development, circadian rhythmicity and mushroom body function (Zhao *et al.*, 2005; Fogg *et al.*, 2014). An interaction between D-MEF2 and FAS2 (the fly ortholog of NCAM) in cell-cell communication or adhesion is suggested by our finding that *D-mef2* hypomorphs exhibit an ectopic veination phenotype similar to that reported for *fas2* loss of function mutant clones (Mao & Freeman, 2009). This is supported by experiments on clock neurons where FAS2 and D-MEF2 work together to control their circadian fasciculation and defasciculation that in turn regulate motor output (Blanchard *et al.*, 2010; Sivachenko *et al.*, 2013). A function for D-MEF2 in defasciculation raises a possible parallel to MEF2’s role in synapse elimination in cultured mouse neurons (Flavell *et al.*, 2006). FOXP (forkhead box transcription factors) proteins are also known to function in synapse elimination. Mammalian FOXP2 has been shown to co-localize with MEF2C early in development but to subsequently suppress MEF2C expression in the striatum (Chen *et al.*, 2016), a dopamine rich forebrain region that is important for motor learning and that has cellular organization that been directly compared to the mushroom bodies (Strausfeld & Herth, 2013; Crittenden & Graybiel, 2017). These results are consistent with distinct cellular functions for MEF2 in development, and later in learning. Disruption of FOXP in the α/β mushroom body neurons results in motor problems and delayed decision-making in an associative olfactory-discrimination task (DasGupta *et al.*, 2014; Lawton *et al.*, 2014) but whether this involves D-MEF2 remains untested.

### *D-mef2* function in mushroom body formation

Deletion of murine *mef2* family members impairs normal development of lymphocytes, bone, endothelial cells, photoreceptor cells and neurons (Mao *et al.*, 1999; Potthoff & Olson, 2007; Andzelm *et al.*, 2015; Latchney *et al.*, 2015). MEF2 activity is required for the survival of primary cerebellar granule neurons that are differentiating in culture, as well as P19 embryonal carcinoma cells that have been induced to undergo neurogenesis (Mao & Nadal-Ginard, 1996; Okamoto *et al.*, 2000). The survival of both cell types is dependent upon p38 mitogen-activated protein kinase, which phosphorylates and thereby activates MEF2 (Mao & Nadal-Ginard, 1996; Han *et al.*, 1997; Okamoto *et al.*, 2000). We have now shown that *D-mef2* is essential for the differentiation of mushroom body neurons.

Constitutive loss of *D-mef2* led to a complete or nearly complete failure in mushroom body formation in all of the homozygous *D-mef2* mutant embryos that we examined. In the homozygous lethal line *D-mef2^26-6^*, there was a 15% reduction in DAC-positive mushroom body cells and an 81% reduction of D-MEF2 positive mushroom body neurons. Thus, the hypomorphic mutation in *D-mef2* had a more profound impact on D-MEF2-positive Kenyon cells than on the surrounding D-MEF2-negative/DAC-positive Kenyon cells. We could not detect any mushroom body neuropil in the *D-mef2* null embryos with the immunomarkers anti-DCO and anti-FAS2, indicating either that the remaining DAC-positive Kenyon cells failed to extend processes or that they were too sparse to detect. Modifiers of the phenotype are suggested by the fact that escaper transheterozygous flies showed grossly normal mushroom body morphology as adults. FAS2 mutations were found to disrupt mushroom body development in one study but not in another (Cheng *et al.*, 2001; Kurusu *et al.*, 2002), further highlighting such phenotypic variability in mushroom body development.

We considered three possible explanations for the reduced mushroom body cell number in *D-mef2* mutants. First, the mushroom body neurons may die prematurely. Second, the mushroom body neuroblasts may fail to proliferate normally. Third, the neurons may not differentiate properly, owing either to a fate change or to a block in the differentiation program. To test whether the primary cause of reduced mushroom body cell numbers in *D-mef2* mutants was cell death, we employed the vital dye acridine orange. Acridine orange was applied to homozygous *D-mef2^26-6^* animals at stages of 14, 15, and 16, periods preceding and including the time at which mutants exhibited a clear reduction in the number of mushroom body neurons. At stage 14, a tight cluster of cells in the dorso-posterior brain was stained with acridine orange in both heterozygous and homozygous *D-mef2^26-6^* animals. By stage 15 and 16 this staining had subsided however, leaving fewer labeled cells that were scattered throughout the CNS (not shown). Although we observed acridine orange staining in the muscle cells of *D-mef2^26-6^* homozygous embryos as previously reported (Ranganayakulu *et al.*, 1995), we did not detect an increase in cell death within the brains of the mutants compared to controls. Therefore, we did not find evidence of apoptotic cell death in the mushroom body neurons of *D-mef2* mutants. Nor was the phenotype caused by the failure of neuroblasts to form: all four mushroom body neuroblasts were apparent at stage 17 in *D-mef2^22-21^* animals as determined by counting experiments. Moreover, the neuroblasts did not express *D-mef2* and did incorporate BrdU, although we cannot rule out that BrdU incorporation was slowed. In conclusion, we propose that the reduction in the number of mushroom body neurons in *D-mef2* mutants may best be explained by a failure of these cells to form or differentiate properly, which is consistent with the absence of mushroom body processes and the neuronal markers FAS2 and DCO.

## ACKNOWLEDGEMENTS

We thank S. Ahmed and B. Schroeder for technical assistance. We thank Profs. Olson, Nguyen and Lilly for providing anti-D-MEF2 antisera. We are grateful to Prof. K.-A. Han for critical reading of the manuscript. This work was supported by a predoctoral NIMH grant to J. R. C. and NINDS grant 1R35NS097224 to R. L. D.

